# Restriction of SARS-CoV-2 Replication by Targeting Programmed −1 Ribosomal Frameshifting In Vitro

**DOI:** 10.1101/2020.10.21.349225

**Authors:** Yu Sun, Laura Abriola, Yulia V. Surovtseva, Brett D. Lindenbach, Junjie U. Guo

## Abstract

Translation of open reading frame 1b (ORF1b) in severe acute respiratory syndrome coronavirus 2 (SARS-CoV-2) requires programmed −1 ribosomal frameshifting (−1 PRF) promoted by an RNA pseudoknot. The extent to which SARS-CoV-2 replication may be sensitive to changes in −1 PRF efficiency is currently unknown. Through an unbiased, reporter-based high-throughput compound screen, we identified merafloxacin, a fluoroquinolone antibacterial, as a −1 PRF inhibitor of SARS-CoV-2. Frameshift inhibition by merafloxacin is robust to mutations within the pseudoknot region and is similarly effective on −1 PRF of other beta coronaviruses. Importantly, frameshift inhibition by merafloxacin substantially impedes SARS-CoV-2 replication in Vero E6 cells, thereby providing the proof of principle of targeting −1 PRF as an effective antiviral strategy for SARS-CoV-2.

## INTRODUCTION

Severe acute respiratory syndrome coronavirus 2 (SARS-CoV-2), the etiological agent of COVID-19, belongs to a family of zoonotic human coronaviruses. Upon the entry of SARS-CoV-2 into host cells, the first set of viral proteins being translated from the single-stranded genomic RNA are encoded by the long (>21kb) ORF1ab, which takes up ~2/3 of the entire viral genome (Figure 1A) (Finkel et al., 2020; Kim et al., 2020). The long polyprotein encoded by ORF1ab is subsequently processed into 16 individual nonstructural proteins (nsp) by two proteases, PLpro/nsp3 and 3CLpro/nsp5. The 3′ half of ORF1ab, ORF1b, encodes enzymes critical for viral transcription and replication, including an RNA-dependent RNA polymerase (RdRp/nsp12), an RNA helicase (Hel/nsp13), a proofreading exoribonuclease and N7-guanosine methyltransferase (ExoN/nsp14), an endonuclease (NendoU/nsp15), and a 2’-O-methyltransferase (nsp16). In all coronaviruses, translation of ORF1b requires a −1 PRF event (Plant et al., 2005). When ribosomes arrive at the end of ORF1a, instead of continuing elongation and soon terminating at an adjacent stop codon in the 0 frame, a subset of ribosomes backtrack by one nucleotide and are repositioned in the −1 reading frame before continuing the translation elongation cycle, thereby producing a complete ORF1ab fusion polyprotein. The −1 PRF region often contains two main components: a heptanucleotide slippery sequence (UUUAAAC in SARS-CoV-2) and a downstream stable secondary structure acting as a frameshift-stimulating element (FSE), which is thought to facilitate −1 PRF in part by transiently pausing the incoming ribosome, allowing tRNAs to realign within the slippery sequence. RNA pseudoknots are the most common type of FSEs found in a variety of RNA viruses (Atkins et al., 2016; Namy et al., 2006). In SARS-CoV, SARS-CoV-2, and other beta coronaviruses, a three-stem pseudoknot (Figure 1B) has been proposed to act as the productive conformation (Plant et al., 2005), although conformational dynamics of this region have recently been observed (Huston et al., 2020; Lan et al., 2020; Zhang et al., 2020).

**Figure 1.**
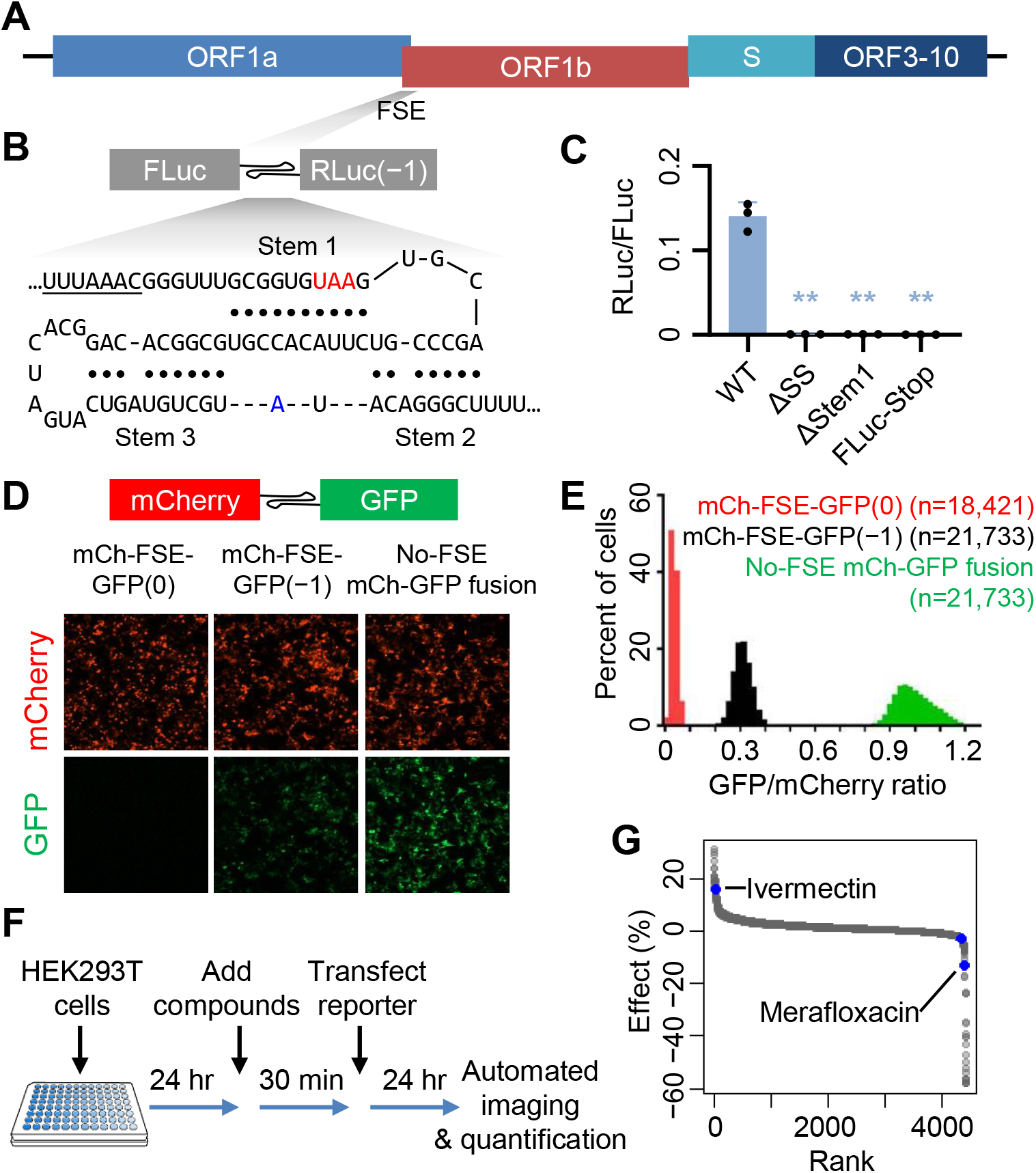
A high-throughput screen identifies SARS-CoV-2 PRF modulators. (**A**) Schematic illustration of the SARS-CoV-2 genome architecture, with the FSE indicated. (**B**) Schematic illustration of the dual luciferase-based −1 PRF reporter design. Watson−Crick base pairs are indicated by filled circles. Each of the three stems in the pseudoknot structure is labeled. The slippery sequence is underlined. The stop codon in the 0 frame is labeled in red. A13533, which varies from a cytosine in SARS-CoV, is labeled in blue. (**C**) Validation of the frameshift reporter. Mutating the slippery sequence (ΔSS), disrupting Stem 1 (ΔStem 1), or adding an in-frame stop codon upstream of pseudoknot eliminates frameshifting. **, p<0.01, two-sided t tests. (**D**) Representative images of cells transfected with mCherry-FSE_CoV-2_-GFP(0), mCherry-FSE_CoV-2_-GFP(−1), or mCherry-GFP. (**E**) Distributions of GFP/Cherry intensity ratios of individual cells transfected with mCherry-FSE_CoV-2_-GFP(−1) (black), mCherry-FSE_CoV-2_-GFP(0) (red), or mCherry-GFP (green). (**F**) Schematic illustration of high-throughput compound screen procedure. (**G**) Ranked effects of 4,434 tested compounds on mCherry-FSE_CoV-2_-GFP(−1) frameshift efficiency. Two validated active compounds, ivermectin and merafloxacin, are labeled in blue.

In contrast to its wide adoption by RNA viruses, −1 PRF is much less prevalent in endogenous mRNAs (Atkins et al., 2016; Clark et al., 2007; Michel et al., 2012). Therefore, viral PRF has been considered an attractive target for specifically interfering with viral gene expression (Brakier-Gingras et al., 2012; Dinman et al., 1998). Indeed, mutations and drugs that alter PRF efficiency have been shown to handicap HIV-1 replication (Brakier-Gingras et al., 2012; Garcia-Miranda et al., 2016; Hung et al., 1998). In addition, an interferon-induced host protein, Shiftless(SHFL), has been shown to interact with HIV-1 FSE RNA, inhibit −1 PRF, and restrict HIV-1 replication (Wang et al., 2019), suggesting that frameshift inhibition has become part of the host antiviral response. Several compounds have recently been shown to modulate −1 PRF of SARS-CoV-2 to varying degrees (Chen et al., 2020; Haniff et al., 2020; Kelly et al., 2020), although the specificity of these compounds and their antiviral activity remain unclear.

Here we show that merafloxacin, a member of a large family of antibacterial compounds known as fluoroquinolones, inhibits −1 PRF of SARS-CoV-2, SARS-CoV, as well as that of other beta coronaviruses. Inhibition of −1 PRF by merafloxacin impeded SARS-CoV-2 replication in Vero E6 cells, indicating that merafloxacin represents a promising lead compound for the development of a novel class of PRF-targeting antivirals.

## RESULTS

### A high-throughput screen identifies SARS-CoV-2 PRF modulators

To quantify −1 PRF efficiency in uninfected cells, we constructed a plasmid-based frameshift reporter (Figure 1B), replacing the stop codon of a firefly luciferase (FLuc) coding sequence (0 frame) with the SARS-CoV-2 FSE (nucleotide 13,460-13,548) including the slippery sequence and the three-stem pseudoknot, followed by a Renilla luciferase (RLuc) coding sequence in the −1 frame. Similar designs have been described in recent studies (Kelly et al., 2020). Several observations confirmed that the relative ratio between RLuc and FLuc activity indeed reported −1 PRF. First, a stop codon is embedded in the −1 frame near the C-terminus of FLuc, ruling out the possibility that RLuc translation initiated within FLuc coding sequence. Second, deleting the slippery site (ΔSS), disrupting Stem 1 of the pseudoknot by a UAC trinucleotide deletion (ΔStem 1), and adding a 0-frame stop codon between FLuc and PRF element (FLuc-Stop), all abolished RLuc activity (Figure 1C), further supporting that RLuc reported on −1 PRF. By using a positive control construct in which FLuc and RLuc were translated continuously without frameshifting, we estimated the PRF efficiency to be ~20% in HEK293T cells, consistent with previous measurements (Kelly et al., 2020).

To make the PRF reporter more suited for high-throughput microscopy screen, we replaced FLuc and RLuc with mCherry and GFP (Figure 1D), respectively. Similar to the luciferase-based reporter, mCh-FSE-GFP(−1) reporter yielded reduced GFP signals compared to the non-frameshifted mCh-GFP fusion construct without an in-frame stop codon (Figure 1D, E). Shifting the GFP back to the 0 frame (mCh-FSE-GFP(0)) completely abolished GFP signal (Figure 1D, E), consistent with the expectation that non-frameshifted ribosomes would terminate at an in-frame UAA stop codon within Stem 1 (Figure 1B).

We treated HEK293T cells in 384-well plates with each of 4,434 compounds at 10 μM final concentration (Figure 1F; **Table S1A**), which included 640 FDA-approved drugs, 1,600 compounds from the Pharmakon 1600 collection, and 1,872 compounds from a Tested-In-Human collection, and transfected cells with the mCherry-FSE-GFP(−1) reporter plasmids. 24 hours after transfection, total cell numbers, as well as mCherry and GFP signals in transfected (mCherry^+^) cells were quantified. GFP/mCherry ratios were compared to the no-PRF mCherry-GFP fusion positive control as well as the mCherry-FSE-GFP(0) negative control. Our high-throughput microscopy screens showed high robustness, with Z′ scores ranging between 0.91 and 0.95.

As expected, the vast majority of the tested compounds had little or no effect on the GFP/mCherry ratio (Figure 1G). Screen actives were selected using mean ± three standard deviations as cutoffs. The majority of top screen actives were false positives due to their intrinsically fluorescence (e.g., doxorubicin (red) and ampiroxicam (green)), which we ruled out by manual inspection of images (**Table S1B**). We repurchased remaining eight candidates and tested each of them using the dual luciferase-based PRF reporter assay. Out of these candidates, two compounds, ivermectin and merafloxacin, were validated as an enhancer and an inhibitor of −1 PRF, respectively (Figure S1). Although ivermectin has been recently shown to have anti-SARS-CoV-2 activity in vitro (Caly et al., 2020) and is currently under clinical trials against COVID-19, it showed significant cytotoxicity concomitantly with −1 PRF enhancement, as indicated by decreased ATP production (Figure S2A) and increased cell death (Figure S2B). In contrast, merafloxacin exhibit modest cytostatic effects at high concentrations (Figure S2C) and did not cause cell death (Figure S2D).

### Merafloxacin specifically inhibits −1 PRF of beta coronaviruses

Merafloxacin, also known as CI-934, belongs to a large group of antibacterial compounds known as fluoroquinolones (Hooper and Jacoby, 2016; Mandell and Neu, 1986). However, none of the 40 other fluoroquinolones in our compound library (Figure 2A) nor additional fluoroquinolones subsequently tested (Figure S3) inhibited −1 PRF, suggesting that the varying moieties at N1, C7, and potentially other positions may be critical for frameshift inhibition (Figure 2B). As expected from the nearly identical FSE sequences of SARS-CoV and SARS-CoV-2, which differ only by one unpaired nucleotide between Stem 2 and Stem 3 (C13533A), merafloxacin inhibited −1 PRF of both coronaviruses with virtually equal efficacy, with IC_50_ of ~20 μM (Figure 2C, D). By comparison, overexpression of SHFL, which has previously been shown to broadly inhibit −1 PRF, only reduced SARS-CoV-2 frameshifting by ~25% (Figure S4), consistent with a recent report (Schmidt et al., 2020).

**Figure 2.**
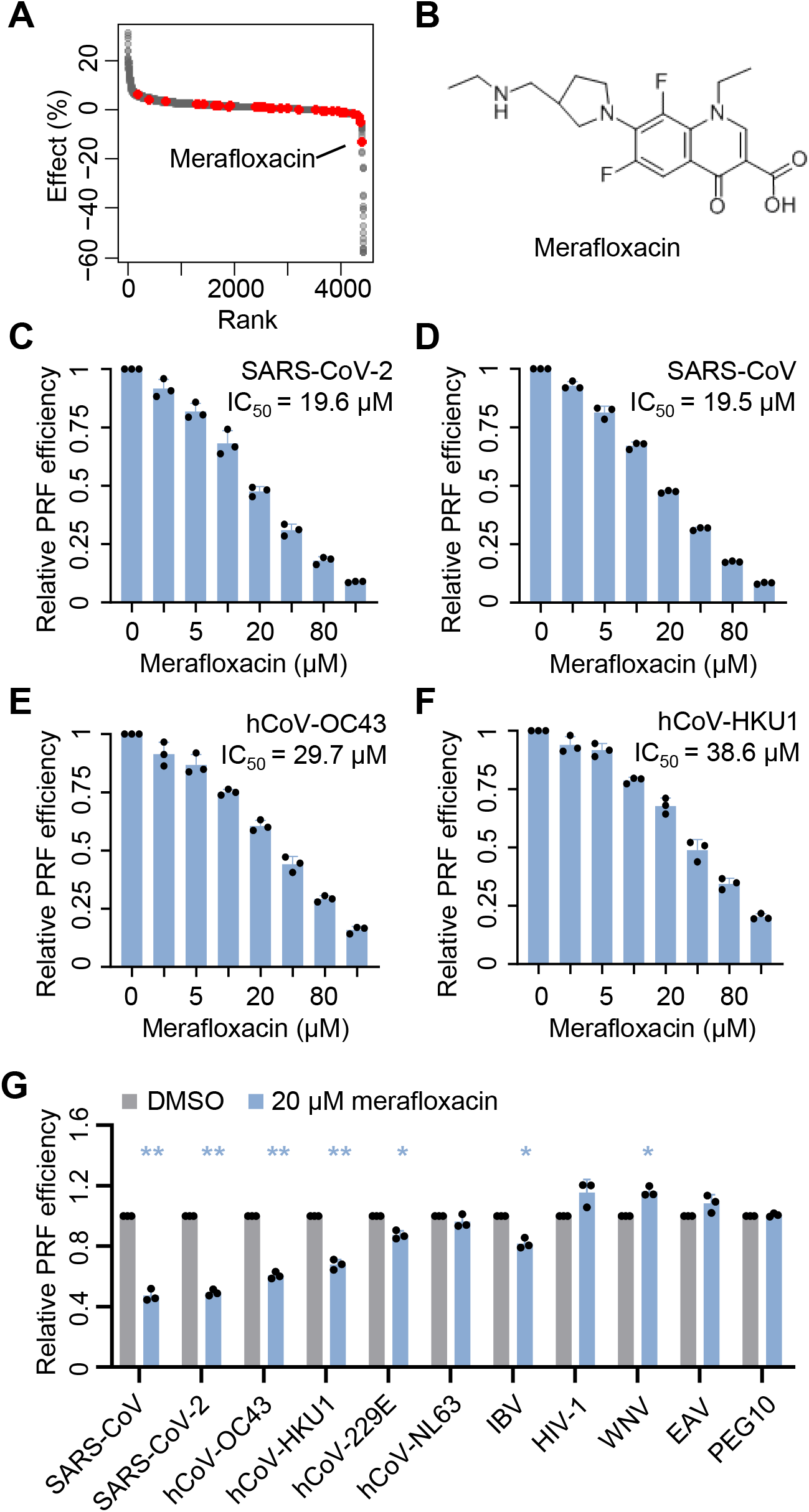
Merafloxacin specifically inhibits −1 PRF of beta coronaviruses. (**A**) Ranked effects of 4,434 tested compounds on mCherry-FSE_CoV-2_-GFP(−1) frameshift efficiency. Fluoroquinolone compounds are labeled in red. (**B**) Chemical structure of merafloxacin. (**C-F**) Dose-dependent inhibition of SARS-CoV-2 (**C**), SARS-CoV (**D**), hCoV-OC43 (**E**), and hCoV-HKU1 (**F**) −1 PRF by merafloxacin, showing IC_50_ values. (**G**) Effects of merafloxacin on −1 PRF of a variety of FSEs. *, p<0.05; **, p<0.01, two-sided paired ratio t tests.

To test whether merafloxacin may inhibit other viral or cellular −1 PRF, we constructed additional reporters using known FSEs from four common human coronaviruses (hCoV-HKU1, hCoV-OC43, hCoV-229E, and hCoV-NL63), avian infectious bronchitis virus (IBV), human immunodeficiency virus 1 (HIV-1), West Nile virus (WNV), equine arteritis virus (EAV), and human PEG10 (paternally expressed gene 10) mRNA. Interestingly, merafloxacin inhibited −1 PRF of other human beta coronaviruses, hCoV-HKU1 and hCoV-OC43, with similar efficacy (Figure 2E, F). In contrast, merafloxacin had much weaker inhibitory effect on −1 PRF of alpha coronaviruses, hCoV-229E and hCoV-NL63 (Figure 2G), whose FSEs have been proposed to form an elaborated pseudoknot structure that substantially differs from that of beta coronaviruses (Herold and Siddell, 1993). Lastly, merafloxacin did no inhibit −1 PRF of HIV-1, WNV, EAV, nor human PEG10 mRNA (Figure 2G). These results suggest that −1 PRF inhibition by merafloxacin is specific to beta coronaviruses, which share a common three-stem pseudoknot architecture.

### Frameshift inhibition by merafloxacin is robust to mutations

A rapidly replicating virus may acquire mutations that confer resistance to an antiviral. To test whether frameshift inhibition by merafloxacin may be escaped by mutations that arise within the FSE, we first introduced mutations that have been documented in the expanding SARS-CoV-2 genome sequence database. Consistent with the essential role of −1 PRF, mutations within the FSE are exceedingly infrequent. Out of 20 documented single-nucleotide substitutions in this region (Hadfield et al., 2018), only four have been observed more than once (Figure 3A). Two of them (C13476U and C13501U) are in Stem 1, each changing a G:C pair to a G:U pair.

**Figure 3.**
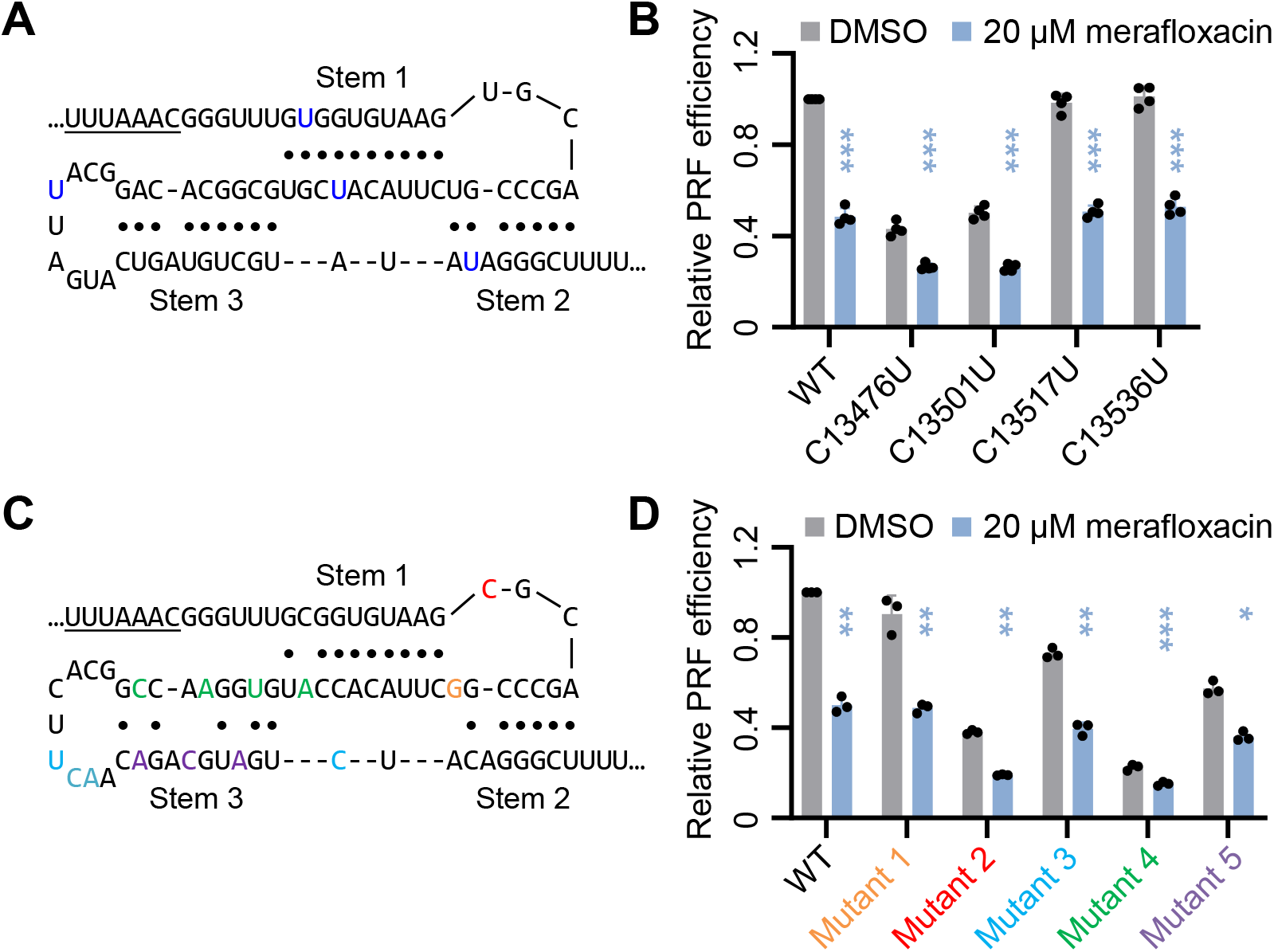
Frameshift inhibition by merafloxacin is robust to mutations. (**A**) Positions of the naturally occurring C-to-U mutations, labeled in blue. (**B**) Effects of merafloxacin on −1 PRF of each FSE natural variant. ***, p<0.001, two-sided ratio t tests. (**C**) Positions of the structure-perturbing, synonymous mutations. Each group of mutations is labeled in the same color. (**D**) Effects of merafloxacin on −1 PRF of each FSE mutant. *, p<0.05; **, p<0.01; ***, p<0.001, two-sided paired ratio t tests.

C13517U is in the terminal loop of Stem 3, whereas C13536U changes a G:C pair in Stem 2 to a G:U pair. Therefore, none of these mutations are expected to disrupt the three-stem pseudoknot architecture. Regardless of the baseline effects of these mutations, merafloxacin inhibited −1 PRF of all of these variants with similar efficacy (Figure 3B).

Having tested these naturally occurring, structure-preserving mutations, we next introduced sets of synonymous mutations intended to perturb the pseudoknot structure (Figure 3C). U13494G (mutant 1, orange) disrupts a basal U:A pair in Stem 2, causing a 10% reduction in frameshifting (Figure 3D). A13519U/G13520C/U13521A/A13533C (mutant 3, cyan) disrupt the palindromic sequence in the terminal loop of Stem 3 (Ishimaru et al., 2013), causing a 27% decrease in frameshifting. G13503A/C13506U/C13509A/A13512C (mutant 4, green) disrupt multiple base pairs in Stems 1 and 3, causing a 77% reduction in frameshifting. U13524A/U13527C/C13530A (mutant 5, purple) disrupt Stem 3, causing a 42% decrease in frameshifting. Unexpectedly, the U13485C mutation (mutant 2, red) within the loop of Stem 1 caused a 62% decrease in frameshifting (Figure 3D), suggesting an uncharacterized function of this presumably unpaired uridine in promoting −1 PRF. Indeed, U13485 and the subsequent G13486 are invariable among all human coronaviruses (Gussow et al., 2020). Despite the wide range of effects of these structure-perturbing mutations, frameshift inhibition by merafloxacin was virtually unchanged (Figure 3D), suggesting that −1 PRF inhibition of merafloxacin is robust to perturbations to the FSE sequence or structure.

### Frameshift inhibition impedes SARS-CoV-2 replication

The identification of a −1 PRF inhibitor allowed us to quantify the effect of frameshift inhibition on viral replication. After infecting Vero E6 cells with SARS-CoV-2, we treated cells with varying concentrations of merafloxacin, and quantified viral yields with plaque formation assays. We found that concomitantly with its inhibition on −1 PRF, merafloxacin inhibited SARS-CoV-2 replication starting from the lowest tested concentration (1.25 μM) (Figure 4A), with an EC_50_ of 2.6 μM and an EC_90_ of 12 μM (Figure 4B). At above 40 μM merafloxacin, viral titer dropped below detection limit (Figure 4A). Importantly, correlating viral titer measurements with the effect of merafloxacin on −1 PRF efficiency (Figure 2C) revealed a near-exponential decrease in viral yield as −1 PRF was increasingly inhibited (Figure 4C), suggesting that −1 PRF efficiency sets the limit of SARS-CoV-2 growth rate.

**Figure 4.**
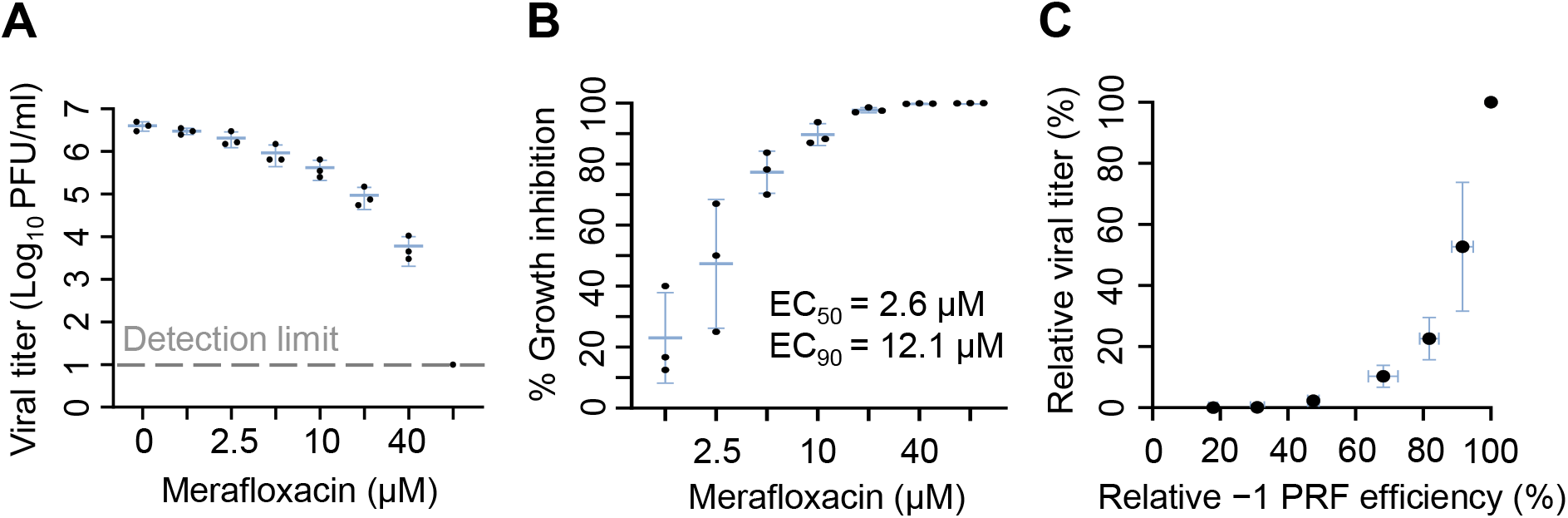
Merafloxacin inhibits SARS-CoV-2 replication in Vero E6 cells. (**A**) Viral yields after 48 hours of merafloxacin treatment. PFU, plaque forming unit. Detection limit is indicated by dashed line. (**B**) Effects of merafloxacin on relative viral growth, based on data from (**A**). EC_50_ and EC_90_ values are shown. (**C**) Relationship between viral titers and relative −1 PRF efficiency, using data from (**A**) and Figure 2C.

## DISCUSSION

Coronaviruses have some of the most efficient FSEs. However, the extent to which SARS-CoV-2 can tolerate changes in −1 PRF efficiency has been unclear. In one scenario, high −1 PRF efficiency could allow coronaviruses to tolerate a small decrease in PRF efficiency while still producing sufficient quantities of replicase components encoded by ORF1b. In this scenario, a large reduction in −1 PRF would be required to inhibit viral growth, and drugs targeting −1 PRF would be less effective for coronaviruses than for viruses with lower −1 PRF efficiency (e.g., HIV-1). The identification of a frameshift inhibitor allowed us to, for the first time, delineate the relationship between frameshift efficiency and viral growth. Our results strongly argue against this scenario. Instead, the near-exponential relationship between viral titer and −1 PRF efficiency (Figure 4C) points to a simple model in which ORF1b translation sets the limit of SARS-CoV-2 growth rate, thereby providing strong support for targeting −1 PRF as an effective antiviral strategy for SARS-CoV-2 and possibly other RNA viruses with high PRF efficiency.

Other non-exclusive models might also explain the high sensitivity of SARS-CoV-2 replication to −1 PRF inhibition. For instance, coronaviruses may require high PRF efficiency to achieve an optimal stoichiometry between the replicase-transcriptase components, which may be disrupted by even a small reduction in −1 PRF efficiency. In this case, increasing PRF efficiency would also be detrimental to the virus, as has been shown for HIV-1 mutants (Garcia-Miranda et al., 2016). Furthermore, multiple ORF1b-encoded replicase-transcriptase components may be rate-limiting for replication, in which case small reductions in each of them would multiply and collectively cause a large effect.

The molecular mechanism by which merafloxacin inhibits −1 PRF is currently unclear. The simplest model would involve the direct binding between merafloxacin and the FSE. Such an interaction may either destabilize the pseudoknot conformation, thereby reducing ribosome pausing and subsequent frameshifting or, on the contrary, further stabilize the pseudoknot structure, thereby causing prolonged stalling, collisions, and/or queuing of the incoming ribosomes. Contrasting ribosome occupancy profiles of either the −1 PRF reporter RNA or the viral genomic RNA with and without merafloxacin treatment may distinguish between these scenarios.

A second possibility is that merafloxacin might stabilize an alternative and unproductive (i.e., non-frameshift-stimulating) FSE conformation. At least one alternative structure with two nested stems has been shown to form in SARS-CoV-2-infected cells by a recent study using dimethyl sulfate probing (Lan et al., 2020). Notably, our ΔStem1 mutant is fully compatible with this alternative conformation, yet it lost all frameshift-stimulating activity (Figure 1C), consistent with the notion that this stem-loop structure is an unproductive conformation. A separate study using in-cell SHAPE probing did not detect this stem-loop structure, but instead observed conformational flexibility of Stem 3 in the three-stem pseudoknot (Huston et al., 2020). Therefore, merafloxacin could plausibly interact and stabilize one or more of these alternative structures, thereby decreasing the fraction of RNAs adopting the productive FSE conformation.

Lastly, merafloxacin might target one or more host factors that mediate or modulate −1 PRF. Although such factors have not been systematically identified, cellular RNA helicases would presumably help unfold the pseudoknot after the ribosome has shifted to the −1 frame and before it continues to translate ORF1b. Considering that the known targets of fluoroquinolones are bacterial DNA topoisomerases (Hooper and Jacoby, 2016), it will be interesting to determine whether their metazoan analogs, some of which have been shown to act on RNAs (Ahmad et al., 2016), may be targeted by merafloxacin.

## METHODS

### Dual-luciferase PRF reporter assay

HeLa and HEK293T cells were cultured in DMEM with 10% fetal bovine serum (ThermoFisher). PRF reporter plasmid DNAs were transfected using Lipofectamine 2000 (ThermoFisher) according to the manufacturers’ instructions. 24 hours after transfection, cells were washed once with phosphate-buffered saline (PBS), and lysed in Glo Lysis Buffer (Promega) at room temperature for 5 min. 1 μL of lysate was diluted with 39 μL PBS before being mixed with 40 μL Dual-Glo FLuc substrate (Promega). After 10 min, FLuc activity was measured in a GloMax 20/20 luminometer (Promega). Subsequently, 40 μL Dual-Glo Stop & Glo reagent was added to the mixture, incubated for 10 min, and measured for RLuc luminescence. The ratio between RLuc and FLuc activities was calculated as frameshift efficiency.

### High-throughput compound screen

HEK293T cells were plated in 384 well plates at the density of 5,000 cells/well. The next day, screened compounds (20 nl of 10 mM stock in DMSO) were added to 20 μl cells using ECHO acoustic dispenser (Labcyte), resulting in 10 μM compound and 0.1% DMSO final concentrations. Cells were treated with candidate compounds for 30 min before transfection of 15 ng mCherry-FSE-GFP(−1) plasmid DNA in each well. 24 hours after transfection, cell nuclei are stained with Hoechst dye. Cell nuclei, mCherry, and GFP signals are imaged using an automated fluorescent microscope (InCell 2200, GE) with a 20x objective, and the acquired images are quantified using the CellProfiler image analysis package. Cell nuclei numbers were quantified as the metric for cell viability. Mean RFP intensity values were first measured in all cells to identify transfected (mCherry-positive) cells, and GFP/mCherry mean intensity ratio was quantified in transfected cells as the metric for PRF efficiency. The no-PRF mCherry-GFP fusion plasmid and the mCherry-FSE-GFP(0) plasmid were used as positive and negative controls for elevated and reduced GFP expression levels, respectively. Screen actives were selected using mean ± three standard deviations as cutoffs.

A total of 4,434 compounds were screened (**Table S1**), including 640 compounds from the FDA-approved library (ENZO), 1,600 compounds from Pharmakon collection (Microsource), and 1,872 compounds from the Tested-In-Humans collection (Yale Center for Molecular Discovery).

### Western blotting

Transfected HEK293T cells were lysed in RIPA buffer on ice for 10 min. After 10 min centrifugation at 4 °C, 20,000 × g, whole cell lysates were mixed with 4X LDS sample buffer (Invitrogen) and denatured at 95 °C for 5 min. Samples were loaded on a 4–12% Bis-Tris SDS-PAGE gel, run at 200 V for 45 min in MOPS buffer, and transferred onto a nitrocellulose membrane (Bio-Rad) in an XCell II Blot module (Invitrogen) (15 V, 45 min). After 1-hour blocking with 5% nonfat dry milk in PBST, the membrane was incubated with primary antibodies (rabbit anti-C19ORF66, Invitrogen # PA5-59815; mouse anti-β-Actin) diluted (1:2000) in 5% milk/PBST at 4 °C with slow shaking overnight. After incubation, membranes were rinsed three times with PBST, and incubated with IR680-or IR800-conjugated secondary antibodies (Li-Cor) diluted (1:10,000) in 5% milk/PBST at room temperature for 1 hour. After three rinses with PBST, membranes were imaged using an Odyssey CLx system (Li-Cor).

### SARS-CoV-2 plaque formation assay

Vero E6 cells (ATCC) were seeded at 2 × 10^5^ cells/well in 12-well plates. The following day, cells were incubated with SARS-CoV-2 viral stock (MOI=5) for 1 hour and washed twice with PBS. Merafloxacin diluted in DMSO was added to the media, mixed briefly, and incubated at 37 °C. After 2 days, media were collected, centrifuged, and supernatants were stored at −80 °C. To quantify viral titers, the collected media were first serially diluted 10-folds with fresh media. 200 μl of each dilution was added to near-confluent Vero E6 cells in 6-well plates and incubated at 37°C for 1 hour with gentle rocking. Subsequently, overlay media (DMEM, 2% FBS, 0.6% Avicel RC-581) was added to each well. After 3 days, cells were fixed with 10% formaldehyde for 30 min, stained with crystal violet for 30 min, and rinsed with deionized water to visualize plaques.

## ACKNOWLEDGEMENTS

We thank Curtis Boswell for help with reagent transportation, Stephen Strittmatter, Pietro De Camilli, and members of the Guo lab for comments on the manuscript. This work was supported by the Yale Scholar in Neuroscience Fund, an NIH Director’s New Innovator Award DP2 GM132930 (to J.U.G.), and NIH awards R01 AI087925 and R01 AI131518 (to B.D.L.). J.U.G. is a NARSAD Young Investigator and a Klingenstein−Simons Fellow in Neuroscience.

## AUTHOR CONTRIBUTIONS

J.U.G. and Y.S. designed the study. L.A. and Y.V.S. optimized and performed the high-throughput microscopy screen and data analysis. Y.S. characterized the active compounds and performed plaque assays. B.D.L. contributed to the study design and trained Y.S. to perform plaque assays. J.U.G. wrote the manuscript with input from all authors.

---

**Figure S1.**
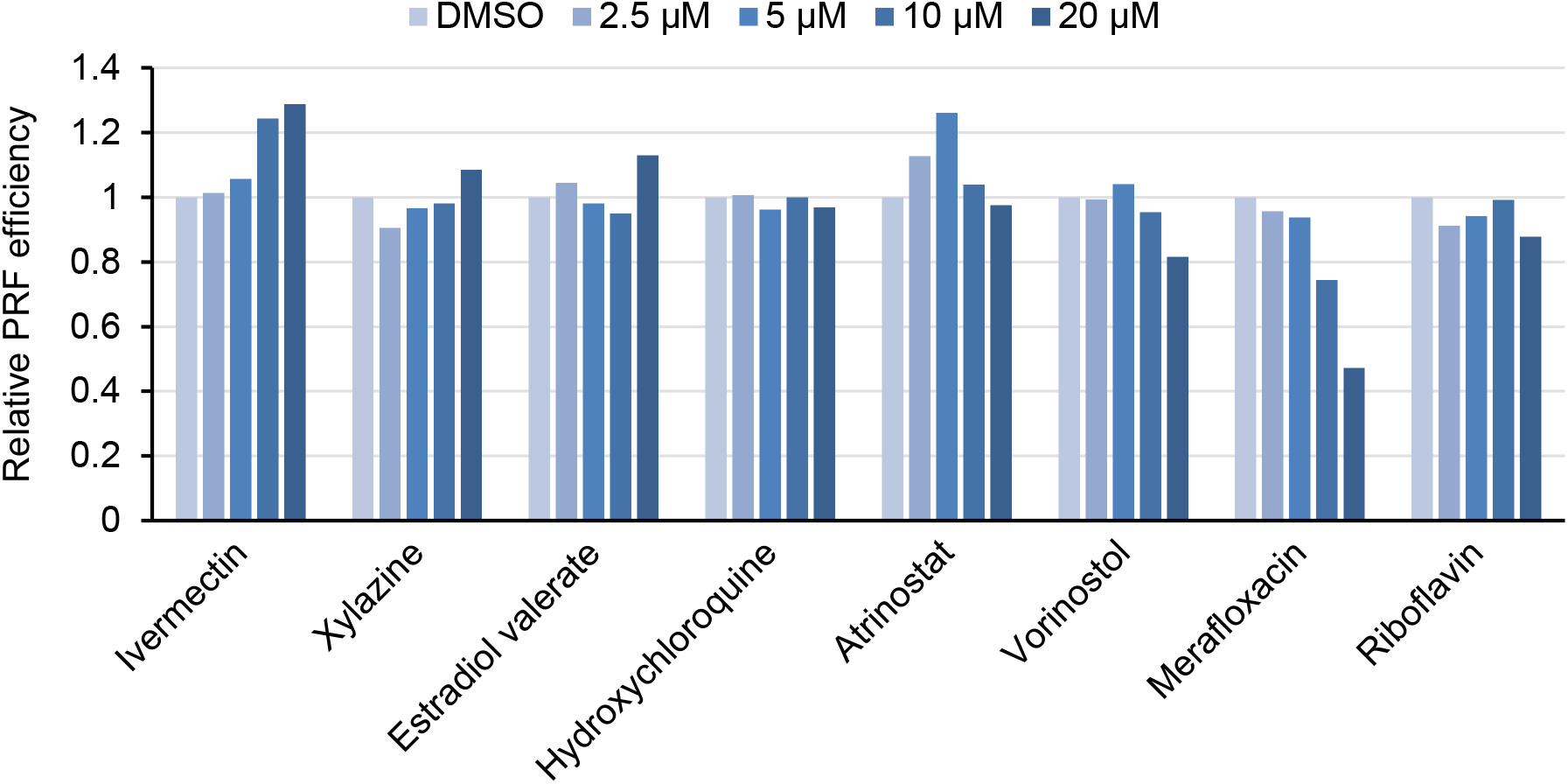
Secondary screen of candidate compounds. Candidate compounds from the microscopy screen were tested using the dual luciferase-based −1 PRF reporter assays.

**Figure S2.**
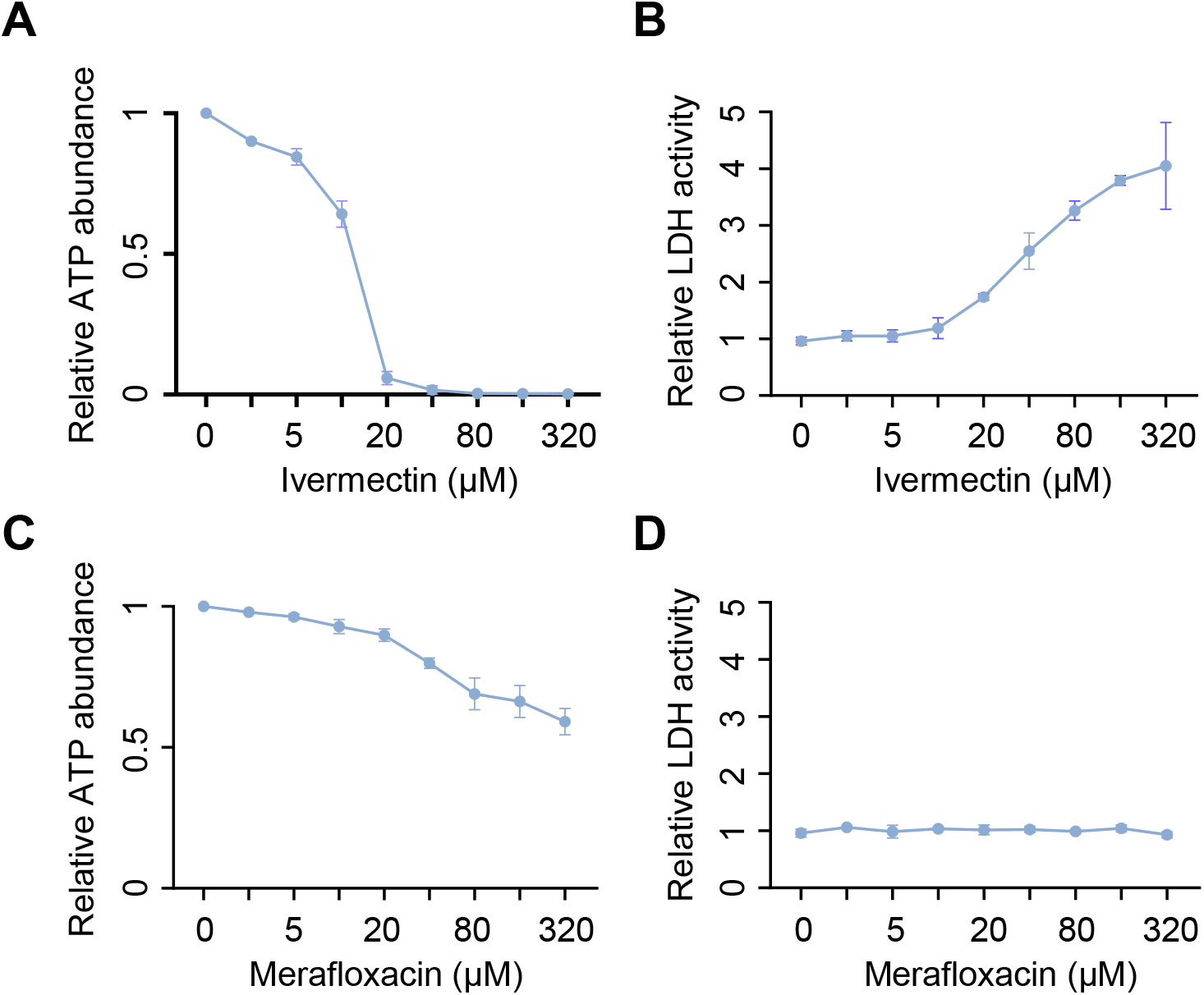
Cytotoxicity of ivermectin and merafloxacin. Cytotoxicity of ivermectin (**A, B**) and merafloxacin (**C, D**) in HeLa cells was quantified by ATP production (**A, C**) and LDH release (**B, D**).

**Figure S3.**
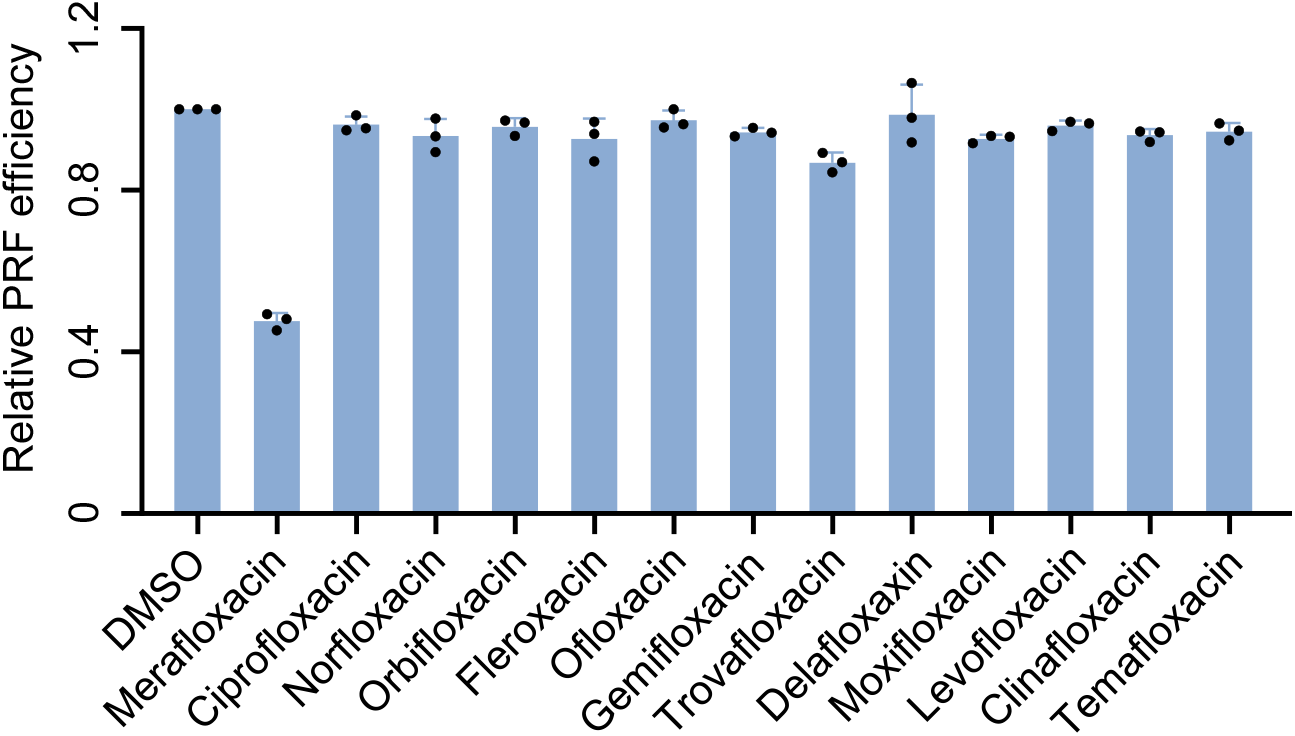
Effect of additional fluoroquinolone compounds on −1 PRF. Each compound was added to a final concentration of 20 μM. Dual luciferase-based −1 PRF reporter assays were used to quantify the effect of each compound.

**Figure S4.**
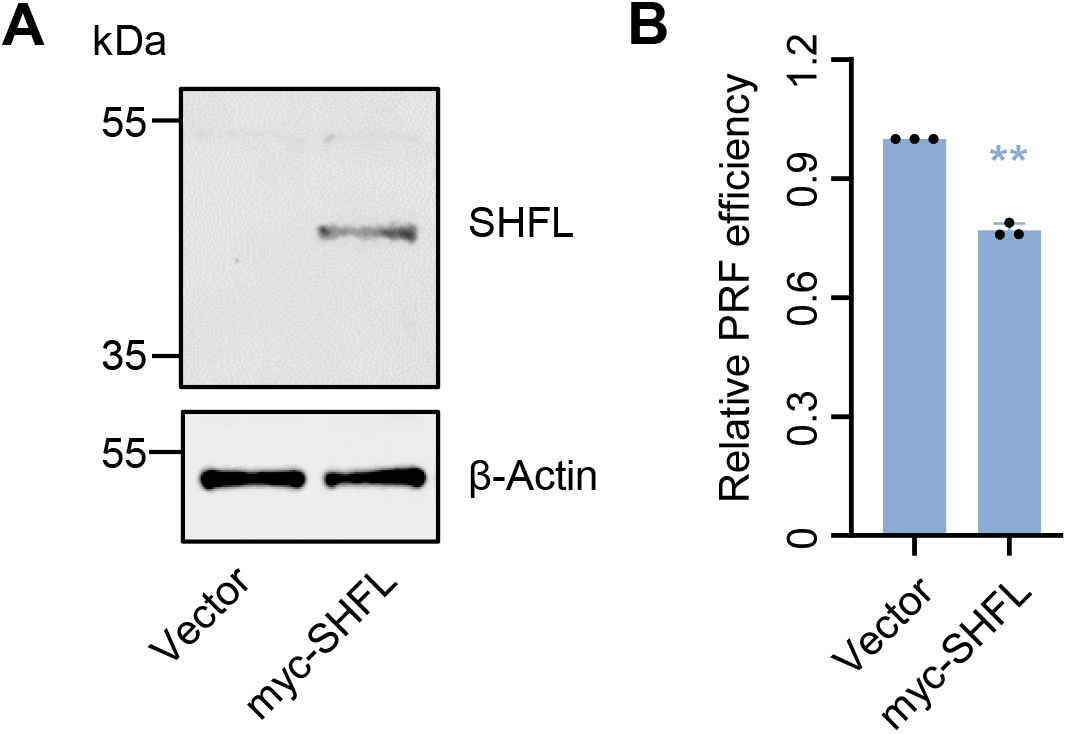
Effect of SHFL overexpression on −1 PRF. (**A**) Western blot showing the overexpression of myc-tagged SHFL in HEK293T cells. (**B**) Effect of myc-SHFL overexpression on −1 PRF efficiency measured with luciferase assays.

## REFERENCES

Ahmad, M., Xue, Y., Lee, S.K., Martindale, J.L., Shen, W., Li, W., Zou, S., Ciaramella, M., Debat, H., Nadal, M., et al. (2016). RNA topoisomerase is prevalent in all domains of life and associates with polyribosomes in animals. Nucleic Acids Res 44, 6335–6349.

Atkins, J.F., Loughran, G., Bhatt, P.R., Firth, A.E., and Baranov, P.V. (2016). Ribosomal frameshifting and transcriptional slippage: From genetic steganography and cryptography to adventitious use. Nucleic Acids Res 44, 7007–7078.

Brakier-Gingras, L., Charbonneau, J., and Butcher, S.E. (2012). Targeting frameshifting in the human immunodeficiency virus. Expert Opin Ther Targets 16, 249–258.

Caly, L., Druce, J.D., Catton, M.G., Jans, D.A., and Wagstaff, K.M. (2020). The FDA-approved drug ivermectin inhibits the replication of SARS-CoV-2 in vitro. Antiviral Res 178, 104787.

Chen, Y., Tao, H., Shen, S., Miao, Z., Li, L., Jia, Y., Zhang, H., Bai, X., and Fu, X. (2020). A drug screening toolkit based on the −1 ribosomal frameshifting of SARS-CoV-2. Heliyon 6, e04793.

Clark, M.B., Janicke, M., Gottesbuhren, U., Kleffmann, T., Legge, M., Poole, E.S., and Tate, W.P. (2007). Mammalian gene PEG10 expresses two reading frames by high efficiency −1 frameshifting in embryonic-associated tissues. J Biol Chem 282, 37359–37369.

Dinman, J.D., Ruiz-Echevarria, M.J., and Peltz, S.W. (1998). Translating old drugs into new treatments: ribosomal frameshifting as a target for antiviral agents. Trends Biotechnol 16, 190–196.

Finkel, Y., Mizrahi, O., Nachshon, A., Weingarten-Gabbay, S., Morgenstern, D., Yahalom-Ronen, Y., Tamir, H., Achdout, H., Stein, D., Israeli, O., et al. (2020). The coding capacity of SARS-CoV-2. Nature.

Garcia-Miranda, P., Becker, J.T., Benner, B.E., Blume, A., Sherer, N.M., and Butcher, S.E. (2016). Stability of HIV Frameshift Site RNA Correlates with Frameshift Efficiency and Decreased Virus Infectivity. J Virol 90, 6906–6917.

Gussow, A.B., Auslander, N., Faure, G., Wolf, Y.I., Zhang, F., and Koonin, E.V. (2020). Genomic determinants of pathogenicity in SARS-CoV-2 and other human coronaviruses. Proc Natl Acad Sci U S A 117, 15193–15199.

Hadfield, J., Megill, C., Bell, S.M., Huddleston, J., Potter, B., Callender, C., Sagulenko, P., Bedford, T., and Neher, R.A. (2018). Nextstrain: real-time tracking of pathogen evolution. Bioinformatics 34, 4121–4123.

Haniff, H.S., Tong, Y., Liu, X., Chen, J.L., Suresh, B.M., Andrews, R.J., Peterson, J.M., O’Leary, C.A., Benhamou, R.I., Moss, W.N., et al. (2020). Targeting the SARS-CoV-2 RNA Genome with Small Molecule Binders and Ribonuclease Targeting Chimera (RIBOTAC) Degraders. ACS Central Science.

Herold, J., and Siddell, S.G. (1993). An ‘elaborated’ pseudoknot is required for high frequency frameshifting during translation of HCV 229E polymerase mRNA. Nucleic Acids Res 21, 5838–5842.

Hooper, D.C., and Jacoby, G.A. (2016). Topoisomerase Inhibitors: Fluoroquinolone Mechanisms of Action and Resistance. Cold Spring Harb Perspect Med 6.

Hung, M., Patel, P., Davis, S., and Green, S.R. (1998). Importance of ribosomal frameshifting for human immunodeficiency virus type 1 particle assembly and replication. J Virol 72, 4819–4824.

Huston, N.C., Wan, H., Araujo Tavares, R.d.C., Wilen, C., and Pyle, A.M. (2020). Comprehensive in-vivo secondary structure of the SARS-CoV-2 genome reveals novel regulatory motifs and mechanisms. bioRxiv.

Ishimaru, D., Plant, E.P., Sims, A.C., Yount, B.L., Jr., Roth, B.M., Eldho, N.V., Perez-Alvarado, G.C., Armbruster, D.W., Baric, R.S., Dinman, J.D., et al. (2013). RNA dimerization plays a role in ribosomal frameshifting of the SARS coronavirus. Nucleic Acids Res 41, 2594–2608.

Kelly, J.A., Olson, A.N., Neupane, K., Munshi, S., San Emeterio, J., Pollack, L., Woodside, M.T., and Dinman, J.D. (2020). Structural and functional conservation of the programmed −1 ribosomal frameshift signal of SARS coronavirus 2 (SARS-CoV-2). J Biol Chem 295, 10741–10748.

Kim, D., Lee, J.Y., Yang, J.S., Kim, J.W., Kim, V.N., and Chang, H. (2020). The Architecture of SARS-CoV-2 Transcriptome. Cell 181, 914–921 e910.

Lan, T.C.T., Allan, M.F., Malsick, L.E., Khandwala, S., Nyeo, S.S.Y., Bathe, M., Griffiths, A., and Rouskin, S. (2020). Structure of the full SARS-CoV-2 RNA genome in infected cells. bioRxiv.

Mandell, W., and Neu, H.C. (1986). In vitro activity of CI-934, a new quinolone, compared with that of other quinolones and other antimicrobial agents. Antimicrob Agents Chemother 29, 852–857.

Michel, A.M., Choudhury, K.R., Firth, A.E., Ingolia, N.T., Atkins, J.F., and Baranov, P.V. (2012). Observation of dually decoded regions of the human genome using ribosome profiling data. Genome Res 22, 2219–2229.

Namy, O., Moran, S.J., Stuart, D.I., Gilbert, R.J., and Brierley, I. (2006). A mechanical explanation of RNA pseudoknot function in programmed ribosomal frameshifting. Nature 441, 244–247.

Plant, E.P., Perez-Alvarado, G.C., Jacobs, J.L., Mukhopadhyay, B., Hennig, M., and Dinman, J.D. (2005). A three-stemmed mRNA pseudoknot in the SARS coronavirus frameshift signal. PLoS Biol 3, e172.

Schmidt, N., Lareau, C.A., Keshishian, H., Melanson, R., Zimmer, M., Kirschner, L., Ade, J., Werner, S., Caliskan, N., Lander, E.S., et al. (2020). A direct RNA-protein interaction atlas of the SARS-CoV-2 RNA in infected human cells. bioRxiv.

Wang, X., Xuan, Y., Han, Y., Ding, X., Ye, K., Yang, F., Gao, P., Goff, S.P., and Gao, G. (2019). Regulation of HIV-1 Gag-Pol Expression by Shiftless, an Inhibitor of Programmed −1 Ribosomal Frameshifting. Cell 176, 625–635 e614.

Zhang, K., Zheludev, I.N., Hagey, R.J., Wu, M.T.-P., Haslecker, R., Hou, Y.J., Kretsch, R., Pintilie, G.D., Rangan, R., Kladwang, W., et al. (2020). Cryo-electron Microscopy and Exploratory Antisense Targeting of the 28-kDa Frameshift Stimulation Element from the SARS-CoV-2 RNA Genome. bioRxiv.

